# The carboxyl-termini of RAN translated GGGGCC nucleotide repeat expansions modulate toxicity in models of ALS/FTD

**DOI:** 10.1101/2020.05.27.119651

**Authors:** Fang He, Brittany N. Flores, Amy Krans, Michelle Frazer, Sam Natla, Sarjina Niraula, Olamide Adefioye, Sami J. Barmada, Peter K. Todd

**Author notes:** Contact, 700 University Blvd, Kingsville, TX 78363, USA, 4005 BSRB, 109 Zina Pitcher Place, Ann Arbor, MI 48109-2200, USA.

## Abstract

An intronic hexanucleotide repeat expansion in *C9ORF72* causes familial and sporadic amyotrophic lateral sclerosis (ALS) and frontotemporal dementia (FTD). This repeat is thought to elicit toxicity through RNA mediated protein sequestration and repeat-associated non-AUG (RAN) translation of dipeptide repeat proteins (DPRs). We generated a series of transgenic *Drosophila* models expressing GGGGCC (G_4_C_2_) repeats either inside of an artificial intron within a GFP reporter or within the 5’ UTR of GFP placed in different downstream reading frames. Expression of 484 intronic repeats elicited minimal alterations in eye morphology, viability, longevity, or larval crawling but did trigger RNA foci formation, consistent with prior reports. In contrast, insertion of repeats into the 5’ UTR elicited differential toxicity that was dependent on the reading frame of GFP relative to the repeat. Greater toxicity correlated with a short and unstructured carboxyl terminus in the glycine-arginine (GR) RAN protein reading frame. This change in C-terminal sequence triggered nuclear accumulation of all three RAN DPRs. A similar differential toxicity and dependence on the GR carboxyl terminus was observed when repeats were expressed in rodent neurons. The presence of the native carboxyl-termini across all three reading frames was partly protective. Taken together, these findings suggest that carboxyl terminal sequences outside of the repeat region may alter the behavior and toxicity of dipeptide repeat proteins derived from GGGGCC repeats.

## Introduction

Nucleotide repeat expansions are a common cause of neurodegenerative disease. Transcribed repetitive sequences can elicit toxicity through at least two possible mechanisms. First, repeats as RNA can bind to and sequester specific proteins and preclude them from performing their normal functions [1]. Alternatively, repeats can be translated into proteins that accumulate and elicit toxicity in model systems [2]. Recently, it was recognized that translation of these repetitive sequences can occur in the absence of an AUG start codon through a process known as repeat associated non-AUG (RAN) translation [3,4]. As such, repeat expansions located outside of protein coding open reading frames can also produce toxic proteins.

In 2011, two groups simultaneously identified GGGGCC repeat expansions as a cause of amyotrophic lateral sclerosis and frontotemporal dementia (C9ALS/FTD) [5,6]. This mutation is common, explaining upwards of 40% of all familial cases of ALS and FTD and 10% of all sporadic cases of these conditions. Most people have only a few repeats, with a cutoff for normal defined as less than 22-28 [7]. Pathologic expansions are usually hundreds to thousands of repeats. The repeat resides the first intron of *C9ORF72*, a gene that is highly expressed in the brain but whose function remains unclear, though evidences indicate the encoded protein may act as guanine exchange factors for activating Rab proteins [8–10]. It is bi-directionally transcribed, such that both G_4_C_2_ and C_4_G_2_ repeat RNAs are produced, and RNA foci from both transcripts are observed in patient-derived cells and tissues [11–13]. Moreover, products of RAN translation are produced from both of these transcripts, leading to 6 different dipeptide repeat (DPR) containing proteins [11,12,14,15]. Repeat expansions also impact *C9ORF72* transcription, isoform choice, and splicing [5,14,16–18].

A series of model systems have provided insights into C9ALS/FTD pathogenesis. *Drosophila* models demonstrate repeat-associated toxicity when the repeat is placed within the 5’ untranslated region (UTR; hereafter referred to as 5’ leader) of a transgene such as GFP [19]. In contrast, long repeats interrupted by stop codons elicit little toxicity [20,21]. Flies expressing dipeptide repeat proteins via AUG initiated translation and independent of G_4_C_2_ repeat sequences are also toxic when expressed in flies and cells in some, but not all reading frames. Specifically, toxicity appears greatest with expression of glycine-arginine and proline-arginine repeat proteins in *Drosophila*, with evidence for a role of glycine-alanine proteins in mammalian neurons and model systems [20–25]. Mouse models using interventricular adenoviral delivery of 66 or 149 G_4_C_2_ repeats in isolation exhibit RNA foci, RAN translation, neurodegeneration, and motor phenotypes [26]. Taken together, these studies do not preclude a role for the repeat RNA in disease toxicity, but suggest that RAN translation directly contributes to neurodegeneration in C9ALS/FTD.

Less attention has thus far been placed on the native sequence context of the repeat. The initial screening of expansion of G_4_C_2_ repeats near ALS loci found that though such repeats are quite common in human genome, the expansion of G_4_C_2_ repeats were only detectable in *C9ORF72* gene but not in other ALS loci genes [27], indicating that the context of the expanded repeats at *C9ORF72* gene are quite unique for expanded repeat toxicity. The sequence context may well be critical to determining the final location of the repeat RNA, its interactions with translational machinery, and the RNA binding proteins with which it interfaces. Tran et al. addressed this issue in *Drosophila* by providing evidence that (a) repeats located within introns are less toxic than repeats placed in 5’m^7^G capped and polyadenylated transcripts, and (b) this enhanced toxicity was associated with increased DPR production [21,28]. However, sequence context may also alter the final translated peptides produced by RAN translation and this could influence their toxicity. For example, in Huntington disease, expression of a *huntingtin* (*HTT*) isoform containing exon 1 alone with the CAG repeat produces a highly toxic and aggregate-prone factor that accumulates in patient brains [29,30]. More recently, Sellier et al. demonstrated a role for the carboxy-terminus of FMRpolyG, a RAN translated protein derived from a CGG repeat expansion in the Fragile X gene, as critical for repeat toxicity in models of Fragile X-associated Tremor Ataxia Syndrome (FXTAS) [31]. We therefore sought to explore whether the repeat location within a transcript, and the sequences 3’ to the repeat that might influence the final RAN translated products, have any influence on GGGGCC repeat toxicity in disease models.

Here we show that expression of up to 484 interrupted G_4_C_2_ repeats within a *Drosophila* intron exhibit minimal toxicity, largely consistent with prior studies [20,21,28]. In contrast, relatively short (28) repeats placed in the 5’ UTR of GFP elicited marked toxicity in *Drosophila* in some, but not all, reading frames relative to a downstream reporter. This differential toxicity correlated with the size and content of the carboxyl terminus of the GR reading frame-with larger carboxyl termini or GFP fusions leading to low toxicity. The relative toxicity observed in these *Drosophila* models correlates with nuclear accumulation of GR DPR-containing proteins and is recapitulated in rodent neurons. The inclusion of the native carboxyl termini provides some neuroprotective effects. Taken together, these findings suggest that repeat context has a significant influence on its toxicity and identifies a new role for the carboxyl termini in modulating DPR toxicity across model systems.

## Materials and Methods

### Construction of mammalian transfection vectors

Constructs for mammalian expression experiments s were generated using by digesting GFP (G_4_C_2_)_71_ pcDNA3.2 (a gift from Christopher Shaw) with XbaI andinserted into pcDNA3.1 containing GFP. One to 2 nucleotides were inserted upstream of the GFP to result in each reading frame. Using NheI and PmeI restriction enzymes, the (G_4_C_2_)_71_ GFP fragment was inserted into pGW for neuronal expression. To generate plasmids with or without the native C-termini, a small fragment was inserted using AscI and MfeI, replacing GFP in pGW. The ATG-V5 fragments were inserted upstream of the repeat using Acc65I and NotI in pGW.

### *Drosophila* Fly Stocks/Genetics

All flies were maintained with standard food and culture conditions at 25^°^C, while all crosses were made and maintained at 29^°^C unless otherwise stated. Fly lines from the Bloomington Stock Center are: GMR-GAL4 (#8605), and UAS-GFP (multiple lines). Actin5C-GAL4/CyO (a gift from Zhe Han), motor neuron specific OK6-GAL4 (a gift from Cathy Collins), RU-486 inducible Tubulin-GAL4 (tub5) driver line (gift from Scott Pletcher).

The sequences for intronic and 5’ leader fly lines are detailed in Supplemental Table S1. Briefly, for intronic repeat lines, a mini intron from the fly *Prospero* gene [32] was inserted into the middle of GFP, synthesized commercially (Genewiz, NJ), and cloned into the EcoRI-XbaI site of pUAST. An intronic fragment with either 3 or 28 G_4_C_2_ repeats along with 40nt of upstream and 260nt of downstream intronic sequence from human *C9ORF72* was PCR amplified according to [33] from genomic DNA and inserted into the intron. The 21 repeat tiling fragment was generated using primer NotI-C9 AnchorR+PspOMI tiling (Supplemental Table 2). The 28 repeat insert was generated using primer NotI-C9R+XhoI-C9F directly from genomic DNA of an ALS patient with 28 repeats. Repeat blocks of (G_4_C_2_)_21_ bracketed by NotI and PspOMI restriction sites were then serially inserted next to this repeat, producing intermediate 49, 70, 91, and 121 repeat containing constructs. This element was then concatamerized to generate lines with 242 (GFP-iC_242_) or 484 (GFP-iC_484_) interrupted repeats.

5’ leader repeat lines were generated by cloning the same 28 repeat sequence described above with approximately 30nt on either side of intronic sequence into the 5’UTR of pEGFP-N1 (Takara Bio USA) in the 0+ reading frame. The repeat and GFP were then subcloned into the NotI site in pUAST. To induce frameshifts, annealing primers were inserted into the AgeI site between the repeat and the AUG of GFP (Supplemental Table S2). pUAST vectors carrying the respective inserts were used to generate transgenic lines by standard p-element insertion (BestGene, CA).

For the fly lines with or without native C-termini, the sequence flanking the (G_4_C_2_)_69_ repeats in constructs for neuron transfection were digested by KpnI and XbaI, and the inserts were cloned to the vector pUAST-AttB (Drosophila Genomics Resource Center, IN) digested with the same set of enzymes. The resulting pUAST-AttB vectors were then used for site-specific transgenesis using PhiC31 integrase technique to locus AttP40 [34] (BestGene, CA). All constructs were verified by Sanger sequencing and agarose gel sizing (Supplemental Figure S1A).

### Eye phenotype imaging and quantification

Representative fly eye images were taken by Leica M125 stereomicroscope and photographed with a Leica DFC425 digital camera as previously described [35]. Eye morphology of 1-2 day post eclosion flies was quantitatively scored as previously described [36]. Briefly, we used the following criteria: supernumerary inter-ommatidial bristles, abnormal bristle orientation, ommatidium fusion, ommatidium pitting, disorganization of ommatidial array, retinal collapse. The presence of each feature will be given 1 point, and additional 2 points were given if more than 5% of the eyes were affected, or 4 points if more than 50% of eyes were affected. Higher score means the eyes are more degenerated. Over 100 flies were scored per genotype in at minimum of three independent crosses. Scores were calculated and are presented as mean ± SEM.

### Fly viability and eclosion rates

Analysis of eclosion rates were performed as described [37]. Each transgenic line was crossed to Actin5C-GAL4 (act5C-GAL4) ubiquitous driver balanced over a marker chromosome (CyO), on standard food at 29^°^C, if the transgene elicited no toxicity, then 50% of progeny should have the CyO marker and 50% express the transgene. Over 100 flies of each genotype were scored over multiple crosses. The relative percent progeny carrying the transgene were expressed as a percent of total eclosed flies. Analysis of viability post eclosion was performed as described [38].

### Larval crawling assay

Transgenic lines were crossed to the motor neuron specific driver line OK6-GAL4 at 29^°^C, and the 3^rd^ instar larvae with desired genotypes were collected. Crawling assay was performed with slight modification from [39]: 3^rd^ instar larvae were collected by adding 20% sucrose on the food, briefly washed with MilliQ water, and then maintained in a water droplet until being analyzed. The larvae were accommodated at room temperature on 2% agarose gel in 10cm petri dish for 1min, and then the crawling distance in the next 1min was recorded and processed by ImageJ. At least 50 larvae from the appropriate genotype were recorded and pooled as the same genotype.

### *Drosophila* Lifespan Assay

The UAS transgenic lines were crossed to Tub5-GAL4 (tub5) Geneswitch driver flies on standard food absent of RU-486 at 29^°^C and adult offspring of the desired genotypes were collected 2-3 days after eclosion and transferred to standard fly food without yeast granules containing 200μM RU-486. The flies were transferred to fresh food with drug every 3-4 days. Each genotype started with at least 4 vials of 25 flies/vial (2 vials of males and 2 vials of females) and the survival was determined daily or every other day for 50 days or until all flies were dead. For each genotype, at least 2 individual lines were examined.

### Generation of specific dipeptide antibodies

The rabbit polyclonal antibodies were generated by Abclonal (Cambridge, MA). Generation of (GA) and (GR) specific antibodies was performed as previously described [40]. To detect GP, we raised antisera against a synthetic (GP)_6_ peptide. All antibodies were affinity purified prior to usage.

### Western Blotting

Protein quantification was as previously described and imaged on standard film [41,42]. Briefly, fly head or cell lysates were lysed in RIPA buffer with protease inhibitors (Roche) and passed through a 27-gauge needle to shear DNA. Equal amounts of protein were run on a 12% SDS polyacrylamide gel. After transfer to PVDF membrane, blots were incubated with the following antibodies: monoclonal mouse anti-GFP (Sigma, MO; clone 7.1 and 13.1, 1:1000), mouse anti-Tubulin (DHSB, IA; clone E7, 1:1,000), mouse anti-V5 (Thermo Fisher, CA; 1:1000), rabbit anti-GA (1:100), rabbit anti-GP (1:5000), rabbit anti-GR (1:5000) or mouse anti-β-Actin (Sigma, MO; 1:5000). At least three independent experiments were performed and scanned films were processed and quantified using ImageJ software.

### In Situ Hybridization

1-3d post eclosion fly heads from each genotype crossed to GMR-GAL4 driver were isolated, immediately frozen in OCT media, and then cryosectioned to 10μm. Transverse sections were then fixed with 4% paraformaldehyde in 1x PBS for 15min, washed 3x in 1x PBS, and permeabilized with 2% acetone for 5min. *In situ* hybridization was performed as follows: sections were prehybridizied with 50% formamide in 2x SSC for 30min in room temperature and then hybridized with 500μl hybridization solution in a light proof box for overnight in 37^°^C. The hybridization solution is as follows: 0.6ng/ml Cy5 labeled 2’-O-Me-(CCCCGG)_5_ RNA probe (IDT DNA Technologies, IA), 0.02% BSA, 132mg/ml yeast RNA (Sigma, MO), 1μl RNAse inhibitor (Sigma), 50% formamide in 2x SSC. After hybridization, slides were washed with 3x 30min of 50% formamide in 0.5x SSC at 56^°^C, followed by 3×10min wash of 0.5x SSC wash at room temperature. After washes, samples were dried for 5min, incubated with 100μl Prolong Gold with DAPI (Thermo Fisher, MA) for 1 hour, and examined on an Olympus FV1000 confocal microscope with identical laser settings for each slide. Images were overlaid and quantitated in ImageJ software.

### Cell based Immunofluorescence

COS-7 cells were maintained 37°C in 5% CO_2_ incubators. Dulbecco’s modified Eagle’s medium (DMEM) supplemented with 10% fetal bovine serum and 1% Pen-Strep was used as culture media. Cells were transfected using Lipofectamine LTX with Plus Reagent (Thermo Fisher, MA) using manufacturer’s protocol. For immunofluorescent detection of GFP, cells were cultured on 4-well chamber slides. 48 hours after transfection, cells were fixed with 4% paraformaldehyde for 15min, washed, permeabilized 0.1% triton X-100, blocked with 5% normal goat serum in 1x PBS containing 0.1% triton X-100, and incubated with GFP and GA (1:100), GP (1:500), or GR (1:500) antibodies overnight. Slides were washed and probed with AlexaFluor 488 labeled goat anti-mouse (Thermo Fisher, MA; 1:500) and Alexa Fluor 555 labeled goat anti-rabbit (Thermo Fisher, MA; 1:500) antibodies and visualized with an Olympus epifluorescence microscope with Slidebook 5.5 software with identical fluorescent settings for each slide.

### RNA isolation and qRT-PCR

Total RNA was extracted from 15 fly heads of each genotype using Trizol (ThermoFisher) and quantitated by Nano-Drop (Thermo Fisher, MA). 1μg of total RNA were then reverse-transcribed to cDNA by iScript cDNA synthesis kit (Bio-Rad, CA). The primers used are detailed in Supplemental Table S2. PCR analysis was performed using the iQ SYBR Green Supermix on a myIQ Single Color qPCR system (BioRad, CA). All runs included a standard dilution curve representing 2x to 0.02x of the RNA concentration utilized for all primer sets to ensure linearity. Equivalent efficiency of individual primer sets was confirmed prior to data analysis. The level of *GFP* was normalized to *RPL32* mRNA for each sample run and expressed as a ratio of levels to GFP-iC3 lines (fold control expression) unless otherwise stated. All samples were run in triplicate in each qPCR run and all data represent at least three independent experiments.

### Primary neuron cultures & transfection

All mammalian rodent work was approved by the Institutional Animal Care and Use Committee (IACUC) at the University of Michigan. Primary mixed cortical neurons were dissected from embryonic day 19-20 Long-Evans rat pups and cultured at 6.0×10^5^ cells/ml in 96 well cell culture plates as previously described [43–45]. Brains of a single litter were combined to maximize cell counts, resulting in neurons from a mixed population of male and female pups. Neurons were cultured in NEUMO photostable medium containing SOS supplement (Cell Guidance Systems, MO) at 37°C in 5% CO_2_. After 4 days in culture, neurons were transfected with 0.2μg DNA (total) and 0.5μL Lipofectamine 2000 (Thermo Fisher, CA) per well in a 96-well plate, as described previously [45]. Neurons were incubated with the DNA/Lipofectamine 2000 mix for 20min at 37°C before rinsing. The remainder of the protocol followed the manufacturer’s protocol.

### Longitudinal fluorescence microscopy

Automated longitudinal fluorescence microscopy began 24 hours post-transfection for 10 days, as previously described [43–45]. Images were acquired using an inverted microscope (Nikon Instruments, NY) with a 20x objective lens, a PerfectFocus system, a Lambda XL Xenon lamp (Sutter Instruments, CA) with 5mm liquid light guide (Sutter Instruments, CA), and either an Andor iXon3 897 EMCCD camera or Andor Zyla4.2 (+) sCMOS camera. All stage, shutter, and filter wheel movements were done using a custom code written in μManager, ImageJ [45].

### Statistical Analysis

For most fly comparisons, scores were analyzed using a non-parametric Kruskal-Wallis ANOVA with Dunn’s correction for multiple comparisons. A Log-rank (Mantel-Cox) Chi square test was performed for *Drosophila* survival rates. Error bars represent the standard error of the mean except for proportion numbers, where the error bars represent the 95% confidence interval.

## Results

### Generation of an intronic repeat model of C9ALS/FTD in *Drosophila*

To create an intronic repeat model of C9ALS/FTD, we inserted the first intron of the fly *Prospero* gene [32] into the sequence of GFP in a fashion that retained the canonical splice site donor and acceptor sequences (Figure 1A). We then inserted either three G_4_C_2_ repeats or concatamers of 28 and 21 G_4_C_2_ repeat units with short interruptions into the intron, up to a maximal repeat length of 484 repeats (Supplemental Figure S1, Supplemental Table S1). After initial studies demonstrated no significant phenotype associated with 28 or 49 intronic repeats (not shown), we focused our analysis on lines expressing 3 (iC_3_), 242 (iC_242_), or 484 (iC_484_) intronic repeats.

**Figure 1:**
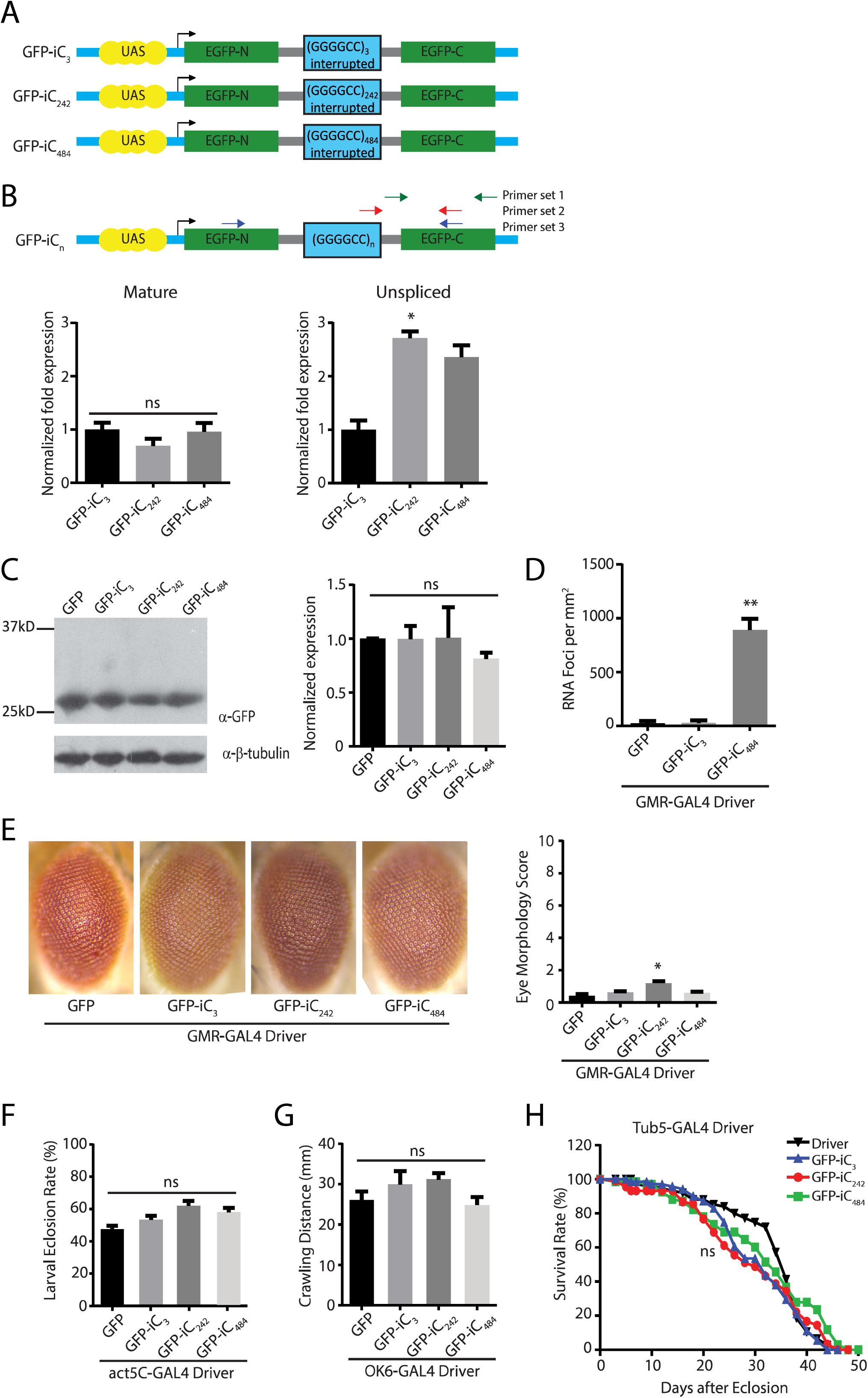
Intronic G_4_C_2_ repeats influence splicing and elicit RNA foci in *Drosophila*. A) Schematic of constructs used to generate transgenic fly lines. The Prospero fly intron 1 (gray lines) was introduced into UAS-GFP (blue lines). Either three G_4_C_2_ repeats or serial (G_4_C_2_)_21-28_ repeat units separated by 14nt interruptions were serially inserted into the intron. B) Locations of the primer pairs used for measuring unspliced, spliced, and total mRNA are show in the schematic. Quantification of the expression of the mature (left) and unspliced (right) GFP product in the indicated fly lines (right). C) Western blot of lysates from heads of G_4_C_2_ intronic repeat flies (left) with quantification of GFP normalized to beta-tubulin (right). D) Quantification of RNA foci in the retina of the indicated intronic fly. E) Representative eye phenotypes from flies of the indicated genotypes crossed to gmr-GAL4 to drive expression in developing ommatidia (left) and quantification of eye phenotype scores (right). F) Quantification of the number of progeny carrying the transgene after flies of the indicated genotypes were crossed to a ubiquitous driver (act5c-GAL4). G) Quantification of the distance crawled by 3^rd^ instar larvae from the crosses of the indicated fly genotypes to a larval motor neuron specific driver (ok6-GAL4). H) Flies carrying the indicated transgenes were crossed to a Tubulin Geneswitch driver (tub5) to allow ubiquitous expression post eclosion. Adult male flies were placed on food containing RU-486 to activate gene expression and their viability was tracked over 50 days. Graphs represent means ± SEM. * p<0.05; ** p<0.01 by Kruskal-Wallis after Dunn’s correction for multiple comparisons.

The G_4_C_2_ repeat as RNA can form a strong G-quadruplex secondary structure that might interfere with RNA metabolism and splicing [18,46,47]. To determine if the repeat taken out of its native context is capable of eliciting alterations in mRNA splicing, we measured the total, unspliced, and spliced mRNA from fly lines containing 3, 242, or 484 intronic repeats (Figure 1B). There was no significant difference in the abundance of spliced *GFP* RNA across repeat sizes (Figure 1B). Consistent with this, the amount of GFP protein expressed in the intronic repeat flies was comparable to that seen in flies expressing *GFP* alone and did not decline with larger repeat sizes (Figure 1C). In contrast, there was a significant repeat-length dependent increase in unspliced pre-mRNA containing the G_4_C_2_ repeat when normalized to *RPL32* (Figure 1B). Additionally, RNA foci were observed in flies expressing long G_4_C_2_ repeats but not short repeats (Figure 1D).

We utilized the UAS-GAL4 system to target expression of the intronic repeats to different tissues. Expression of GFP-iC_3_, GFP-iC_242_, or GFP-iC_484_ in the eye using a GMR-GAL4 driver elicited modest toxicity in GFP-iC_242_ eye phenotypes compared to a control line expressing GFP alone using a standardized toxicity scale [35] (Figure 1E). No significant eye phenotypes were observed in GFP-iC_484_ expressing lines. Similarly, activating ubiquitous expression during development using an Actin5C-GAL4 driver elicited no differences in eclosion rates between expanded intron lines and the control GFP line (Figure 1F). We also selectively activated expression of the intronic repeat transgenes in motor neurons using an OK6-GAL4 driver line and evaluated larval crawling as a motor phenotype. We observed no difference between short and long intronic repeats compared to flies expressing *GFP* alone (Figure 1G). To evaluate whether a reduction in lifespan might be elicited in adulthood, we activated transgene expression in adult male flies after eclosion using a Geneswitch-Tubulin-GAL4 driver by addition of RU-486 to the fly food. We observed no significant impact of intronic repeat expression on viability (Figure 1H).

### The role of repeat location in GGGGCC toxicity

To determine if the repeat location within a transcript might impact its toxicity, we inserted the same 28 G_4_C_2_ repeat cassette used to generate the intronic repeats into the 5’ leader of GFP rather than an intron (Figure 2A). No start codons were present between the transcription start site and the repeat, although the ATG of GFP was retained 3’ to the repeat. We introduced small frameshift mutations between the repeat and the open reading frame of GFP so that it acted as a reporter for RAN translation in the 0+ (GA), 1+ (GP), or 2+ (GR) reading frames, respectively.

**Figure 2:**
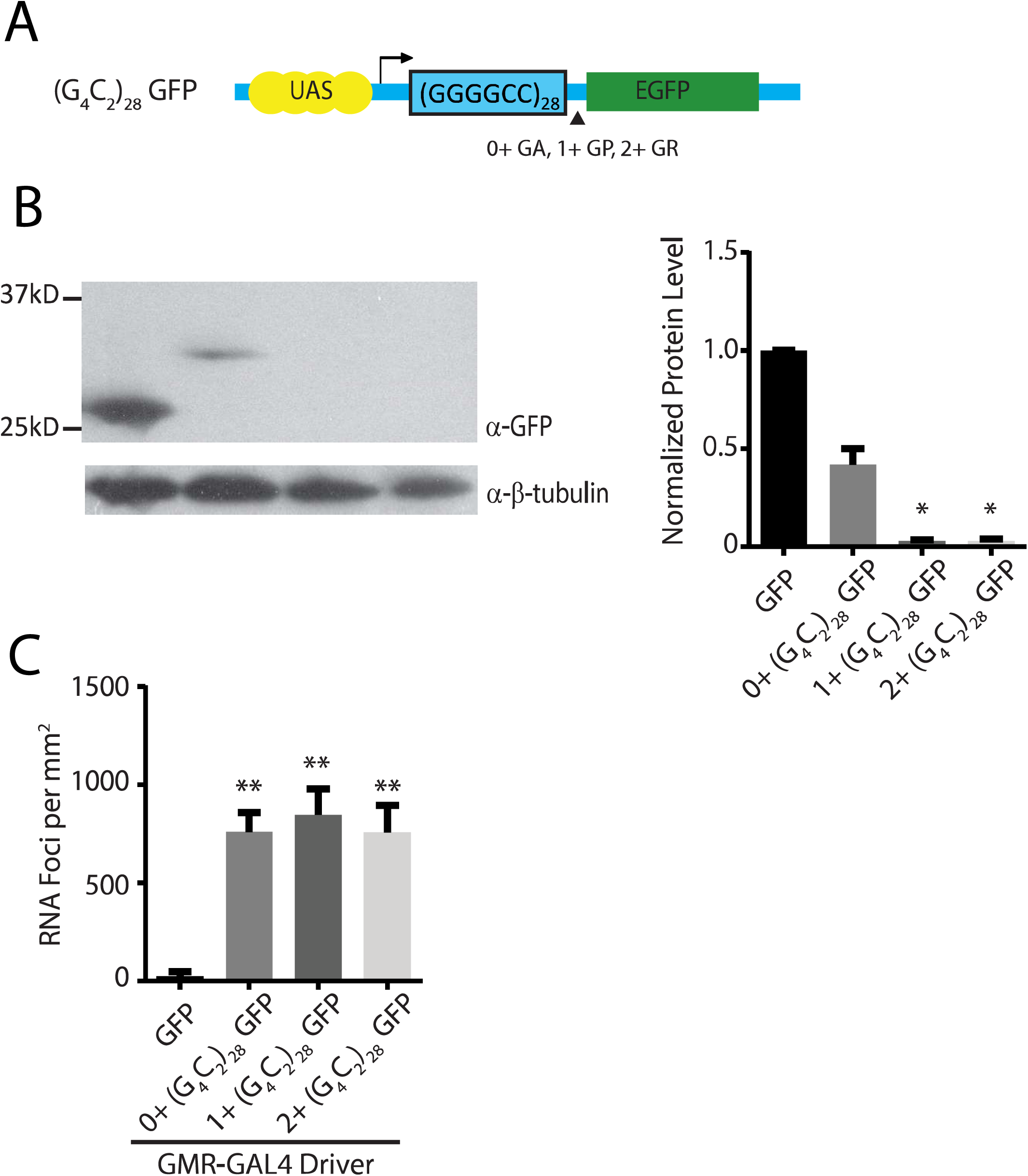
*Drosophila* 5’UTR G_4_C_2_ repeat models. A) Schematic of constructs used to generate transgenic fly lines. (G_4_C_2_)_28_ repeats were inserted into the 5’UTR of GFP without an upstream start codon in all three reading frames relative to the GFP reading frame. B) Western blot (left) of lysates from heads of G_4_C_2_ exonic repeat flies normalized to beta-tubulin (right). C) Quantification of the presence of RNA foci in the retina of the indicated fly lines with. Graphs represent means ± SEM. * p<0.05; ** p<0.01 by Kruskal-Wallis after Dunn’s correction for multiple comparisons.

When the G_4_C_2_ repeat was expressed in an intronic context, levels of GFP protein were similar between short repeats and expanded repeats. However, placing the repeat in the 5’ leader of *GFP* dramatically affected overall *GFP* expression. A higher molecular weight GFP fusion protein product was observed by western blot in the 0+ (GA) frame, but undetectable in the GP (+1) or GR (+2) reading frames (Figure 2B). Expression of all three constructs led to detectable RNA foci (Figure 2C). The presence and abundance of RNA foci did not directly predict toxicity in these flies, similar to prior observations [28].

When expressed in fly ommatidia, 5’ leader repeat fly lines elicited greater toxicity than was observed in the intronic repeat fly lines across a range of measures, consistent with a prior publication [28]. However, surprisingly, there were significant differences in the toxicity observed that was dependent on the reading frame in which the GFP reporter was placed. Specifically, 1+ (G_4_C_2_)_28_ GFP fly lines elicited a marked degeneration of the eye (Figure 3A) and to a lesser extent in 2+ (G_4_C_2_)_28_ GFP flies. Developmental toxicity, as assessed by eclosion rates after ubiquitous expression, and a severe larval motor phenotype observed with isolated motor neuron expression, largely mirrored toxicity findings observed in the eye (Figure 3B-C). Moreover, rapid declines in viability after transgene activation in adulthood were observed in all 5’ leader flies, with the most robust toxicity in 1+ (G_4_C_2_)_28_ GFP fly lines (Figure 3D).

**Figure 3:**
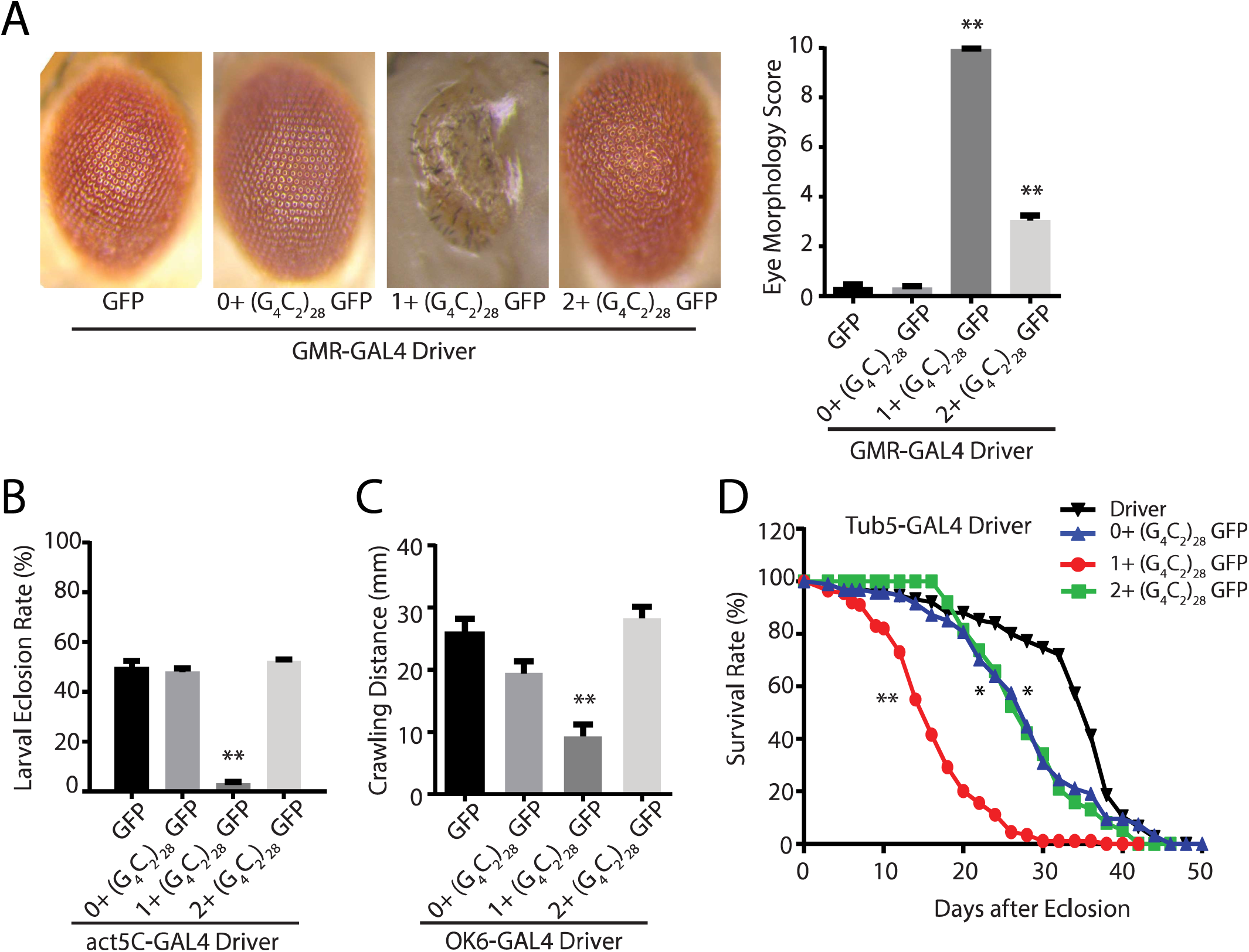
5’UTR G_4_C_2_ repeats with different Carboxyl termini elicit differential toxicity in *Drosophila*. A) Representative eye phenotypes from flies of the indicated genotypes crossed to GMR-GAL4 to drive expression in developing ommatidia (right) and quantification of eye phenotype scores (right). B) The number of progeny carrying the transgene was determined after flies of the indicated genotypes were crossed to a ubiquitous driver (act5c-GAL4). C) Quantification of the distance crawled by 3^rd^ instar larvae from the crosses of the indicated fly genotypes to a larval motor neuron specific driver (ok6-GAL4). D) Flies carrying the indicated transgenes were crossed to a Tubulin Geneswitch driver (tub5) to allow ubiquitous expression post eclosion. Adult male flies were placed on food containing RU-486 to activate gene expression and their viability was tracked over 50 days. Graphs represent means ± SEM. ** p<0.01 by Kruskal-Wallis after Dunn’s correction for multiple comparisons. Log-rank (Mantel-Cox) test for survival, * p<0.05, ** p<0.01.

In an attempt to explain these differences in phenotype, we compared transgene mRNA expression across lines and observed no correlation between toxicity and abundance of the transgene expression (Supplemental Figure S1B). We next analyzed three independent lines for each transgene to control for effects of transgene insertion sites. Phenotypes for different lines of the same transgene were very similar despite significant differences across transgenes (Supplemental Figure S1C). Of note, the DNA sequence 5’ to the repeat in all three transgene constructs were identical, precluding contributions from an upstream initiation event in only one construct as a driving force in the observed phenotypic differences (Supplemental Table S1).

### Accumulation and localization of DPR protein correlates with toxicity

To determine what factors might be triggering the differential toxicity between these *Drosophila* lines, we used a series of new and recently developed [40] antibodies against the three sense strand DPRs: GA, GR, and GP. These polyclonal antibodies recognize epitope tagged versions of the indicated repeat proteins both by western blot and by immunocytochemistry in transfected cells (Supplemental Figure S2A-B). These antibodies also stain perinuclear inclusions in cerebellar slices from ALS patients with *C9ORF72* repeat expansions but not in healthy controls (Supplemental Figure S2C).

We utilized these antibodies to determine the relative abundance and distribution of each DPR in all of the transgenic *Drosophila* lines described. We reasoned that the epitope tags provide information on a single reading frame, but that RAN translation occurs in all 3 reading frames and thus the relative abundance of each product might be informative for their roles in toxicity. Using transverse ommatidial sections derived from *Drosophila* expressing each transgene (UAS-GFP, GFP-iC_3_, GFP-iC_484_, and 0+, 1+, or 2+ (G_4_C_2_)_28_ GFP lines) crossed to GMR-GAL4, we performed immunocytochemistry against each of the three DPRs using these new antibodies. No staining was observed in lines expressing GFP in isolation or in the driver line alone. However, staining was consistently present with both GA and GR antibodies in the intronic fly models, with significantly more staining at larger repeat sizes (Figure 4). In contrast, GP staining was not reliably observed from the intronic fly lines. In 5’ UTR lines there were marked differences in the staining for both GR and GP between lines, with the greatest staining present in the +1 (G_4_C_2_)_28_ GFP lines that exhibited the greatest toxicity (Figure 4C-D). In addition, this same line exhibited a marked increase in both total nuclear staining and in the nuclear-cytoplasmic ratio for all three DPR proteins (Figure 4D), suggesting a correlation between both abundance and cellular localization of these proteins and the observed toxicity in *Drosophila* [20,25,28,48].

**Figure 4:**
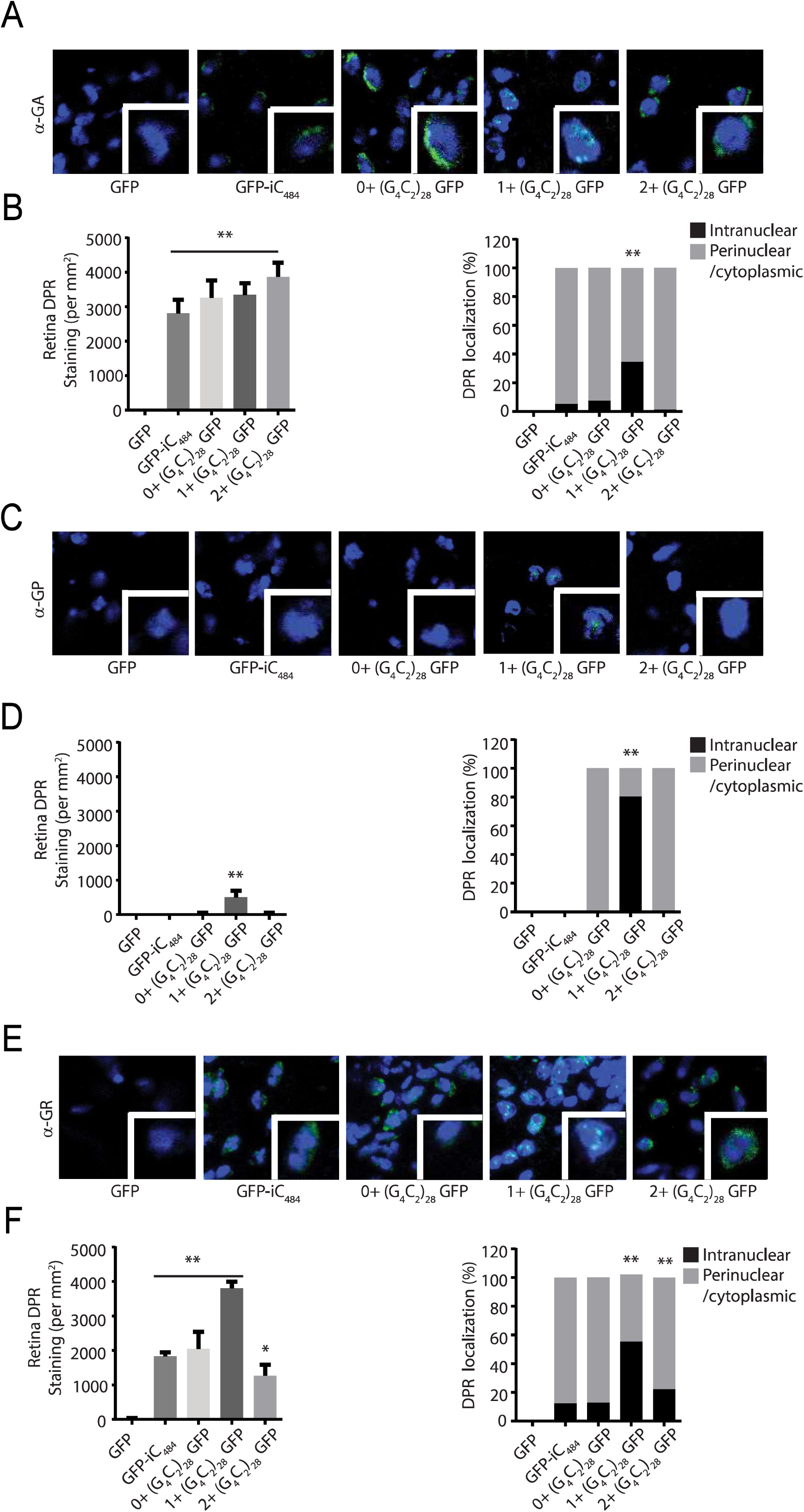
G_4_C_2_ repeats toxicity is associated with nuclear accumulation of RAN translation peptides. A) Representative images of transverse retinal sections from *Drosophila* of the indicated genotypes probed with antibody against GA dipeptide repeats. B) Quantification of total GA repeat staining by immunofluorescence (left) and the percentile of cells with significant nuclear accumulation of GA DRP staining for the indicated genotypes (right). C) Representative Immunofluorescence images of *Drosophila* retinal sections probed with antibodies against the GP dipeptide repeats. D) Quantification of total GP repeat staining by immunofluorescence (left) and the percentile of cells with significant nuclear accumulation of GP DRP staining for the indicated genotypes (right). E) Representative Immunofluorescence image of transverse retinal sections from *Drosophila* of the indicated genotypes probed with antibody against GR dipeptide repeats. F) Quantification of total GR repeat staining by immunofluorescence (left) and the percentile of cells with significant nuclear accumulation of GR DRP staining for the indicated genotypes. Graphs represent means ± SEM. * p<0.05; ** p<0.01 by Kruskal-Wallis after Dunn’s correction for multiple comparisons.

Prior studies have demonstrated that carboxyl terminal sequences and epitope tags can influence the toxicity associated with translated repeat expansions such as polyglutamine proteins [30]. More recently, the carboxyl terminus of the RAN translated protein FMRpolyG that is generated from GGC repeats in FXTAS was found to influence the relative toxicity of this protein through aberrant protein-protein interactions [31]. We therefore analyzed the C-termini of the transgenes used to generate the different exonic flies (Table 1). In the native *C9ORF72* gene, each DPR reading frame has 30-55 amino acids after the repeat prior to the presence of a stop codon. Such C-termini were detectable in ALS patients carrying the expanded G_4_C_2_ repeats according to previous studies [13]. In our exonic flies, 2 reading frames have C-termini over 200 amino acids (one of these frames is GFP) while the third is much shorter (15-21 amino acids).

**Table 1:**
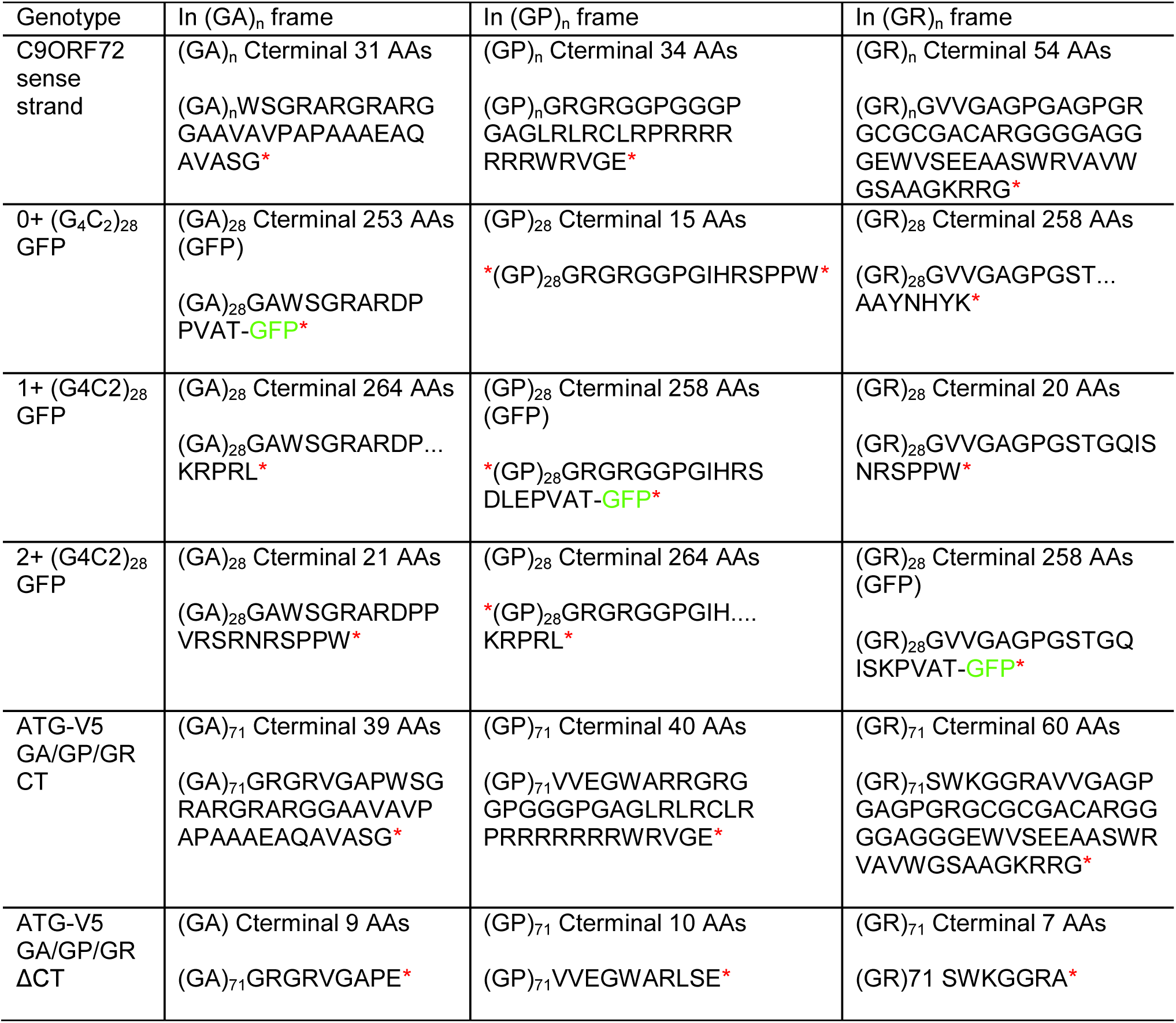
RAN Translation Peptides in G_4_C_2_ repeat transgenic flies and plasmids

Based on these observations, we hypothesized that the shorter C-terminus in the GR reading frame allowed for stabilization of this product within cells that favored its redistribution into the nucleus and a subsequent activation of toxicity. To explore this idea further, we utilized longitudinal microscopy [31,43–45] to track survival of individual neurons expressing sequences similar to those used in the fly experiments (Figure 5A). Primary rodent cortical neurons were transfected with plasmids encoding mApple, to visualize neuronal soma, and each of the 5’ leader G_4_C_2_ constructs. These cells were imaged at 24h intervals over a 10d period, and the time of death for individual neurons determined programmatically using a set of sensitive criteria (rounding or blebbing of the cell body or neurites, loss of fluorescence) validated in previous studies [49,50]. Relative survival in each condition was compared by Cox proportional hazards analysis. As we observed in *Drosophila*, expression of an expanded G_4_C_2_ repeat in the 5’ UTR of GFP was toxic in neurons. The relative toxicity of each construct was again dictated by the reading frame of the GFP tag, with toxicity being greatest in those with 1+ G_4_C_2_ GFP or 2+ G_4_C_2_ GFP, mirroring what was observed in *Drosophila* (Figure 5A).

**Figure 5:**
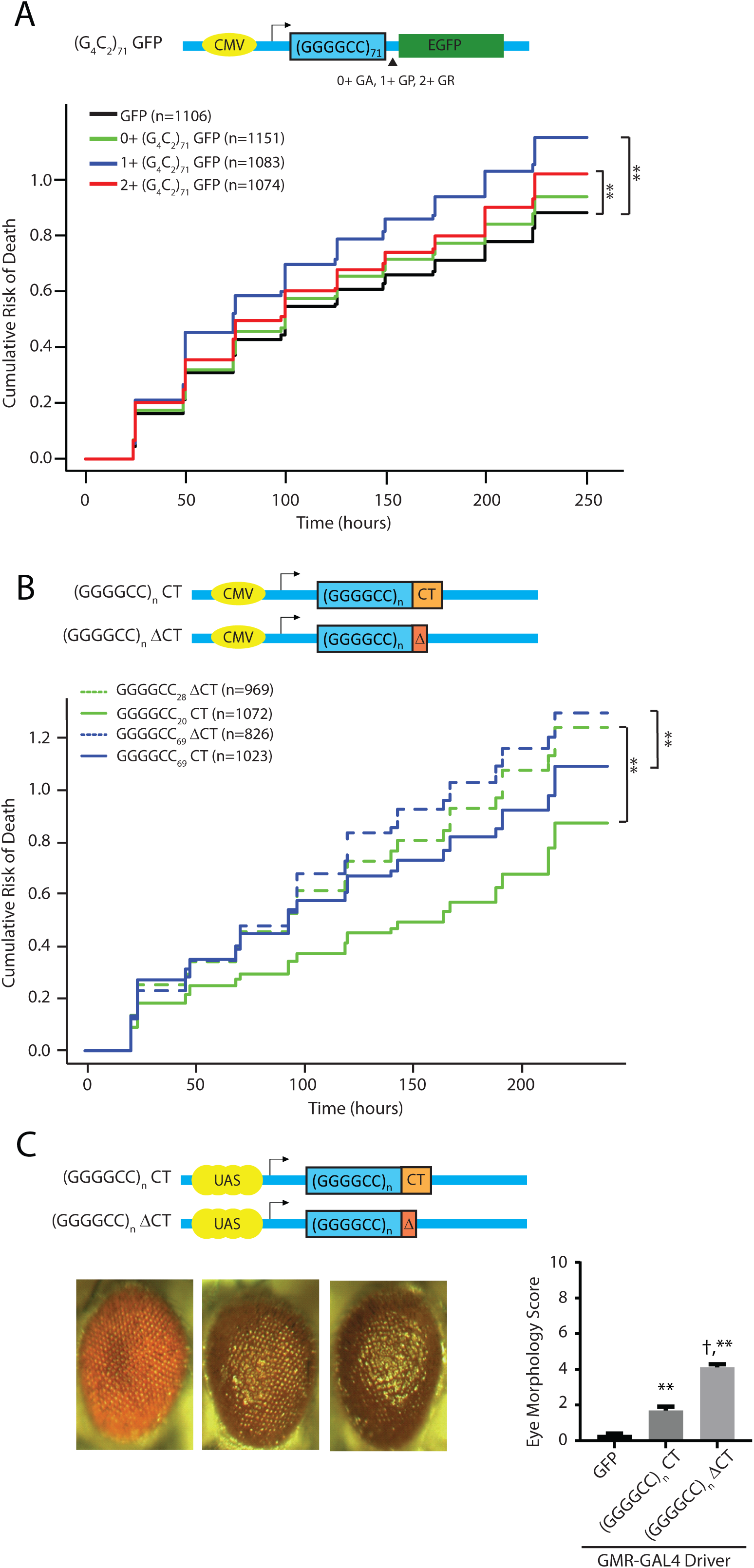
Native Carboxyl terminal sequences reduce GGGGCC repeat toxicity in rodent neurons and in flies. A) Schematic of the plasmids transfected into neurons. Cumulative risk of death in each reading frame when the GGGGCC repeat is in the 5’ leader sequence of GFP. B) Schematic of the plasmids transfected into neurons. Cumulative risk of death of neurons with normal and expanded G_4_C_2_ repeats with and without the native C-termini. ** p<0.01 by Cox proportional hazard analysis. C) Schematic of the plasmids used to generate transgenic fly lines. Representative eye phenotypes from flies of the indicated genotypes crossed to gmr-GAL4 to drive expression in developing ommatidia (left) and quantification of eye phenotype scores (right). Graphs represent means ± SEM.** p<0.01 by Kruskal-Wallis after Dunn’s correction for multiple comparisons compared to control group. † p<0.01 by Kruskal-Wallis after Dunn’s correction for multiple comparisons compared to (G_4_C_2_)_69_ CT group.

To evaluate the potential role of different carboxyl terminal sequences on toxicity, we generated a new series of expression vectors that take into account the length and sequence of the C-termini. One set of vectors have the native C-termini (CT) for all three reading frames. Another set of vectors was generated with shortened C-terminal regions in all three reading frames (ΔCT) (Figure 5B). The G_4_C_2_ repeat with the native C-terminal sequence showed increased toxicity with an increasing repeat size (Figure 5B, solid green vs solid blue line). However, when the native C-termini were removed and replaced with short C-termini, toxicity was significantly increased (Figure 5B, dashed lines vs solid lines). These (G_4_C_2_)_69_ repeats with the native C-termini (CT) or with short C-termini (ΔCT) were also introduced to the same locus (AttP40) in *Drosophila* using pUAST-AttB vectors to achieve equal expression levels. When expressed in the eye, inclusion of the the C-termini partially suppressed the repeat-associated toxicity (Figure 5C).

In an attempt to determine which C-terminal reading frame was most critical for toxicity, we introduced a series of AUG driven V5 tags into the 5’ UTR upstream of the repeat in different reading frames. Enhancing production of GA or GR from these repeats significantly boosted the toxicity of these constructs to a greater degree than placement of an AUG codon in the GP reading frame. However, replacing the native C-terminal sequences with stop codons only enhanced repeat toxicity when the AUG codon was placed in the GR reading frame (Figure 6C-E). Taken together, these data suggest that the native C-termini may mitigate toxicity arising from RAN translation, and in particular toxicity related to GR DPR proteins.

**Figure 6:**
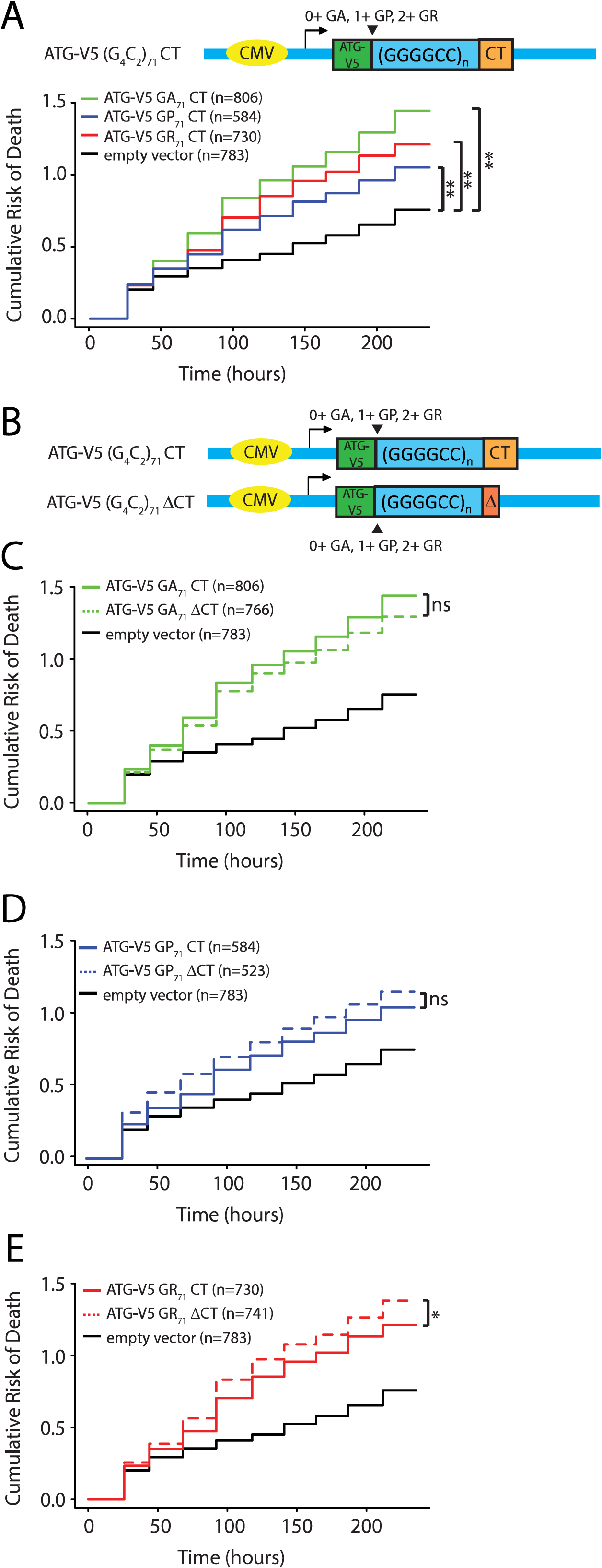
Native Carboxyl terminal sequences decrease GR dipeptide repeat elicited toxicity in rodent neurons. A) Schematic of the plasmid for each individual reading frame with the native C-terminal sequence transfected into neurons with the cumulative risk of death below. B) Schematic of the plasmids for each individual reading frame with different C-terminal sequences transfected into neurons. C) Cumulative risk of death of neurons transfected with GA repeats with expanded repeats with different C termini. D) Cumulative risk of death of neurons transfected with GP repeats with expanded repeats with different C-termini. E) Cumulative risk of death of neurons transfected with GR repeats with expanded repeats with different C-termini. * p<0.05 and** p<0.01 by Cox proportional hazard analysis.

## Discussion

How hexanucleotide repeat expansions in *C9ORF72* elicit neurodegeneration is a topic of intense research [51,52]. Work by multiple groups has established the presence of both nuclear RNA foci and RAN translation derived DPR proteins in tissues and cells from affected patients. Initial studies largely relied on expression of the repeat out of its normal sequence context or production of DPR proteins via AUG-initiated translation. These studies importantly demonstrate that even relatively short RNA repeats and small DPRs can be toxic [19,20,22–26,53–55]. The position of the GGGGCC repeat within an mRNA is a major factor in determining its relative toxicity in *Drosophila*, with repeats residing in an intron exhibiting minimal toxicity while even short repeats in the 5’ leader elicit significant toxicity [28]. In this work we observe that sequences 3’ to the repeat also modulate repeat toxicity by altering the carboxyl termini of DPRs generated via RAN translation. These findings suggest a specific role for C-terminal sequences in modulating repeat toxicity in C9ALS/FTD while making it clear that repeat sequence context needs to be carefully considered in future studies of disease pathogenesis.

Sequences located in coding regions or within the 5’ or 3’ UTR are typically processed to capped and polyadenylated mRNAs and rapidly trafficked out of the nucleus, where they interface with a series of RNA binding proteins and the translational machinery. In contrast, intronic sequences are typically spliced out during transcription, formed into RNA lariats and targeted for debranching and degradation in the nucleus. As such, one potential explanation for the observed differences in toxicity between intronic and 5’ leader sequence repeats is the distribution of the repeat containing transcripts within the cell and their potential for translation. *In vitro* and cell based studies using reporter systems suggest that while GGGGCC repeats can be translated from intronic or bicistronic transcripts, their production is most robust when present in capped and polyadenylated mRNAs [56–60]. Consistent with this prediction, the abundance of specific RAN translation products from relatively short 5’ leader repeats were much greater than that observed from larger intronic repeats (Figures 1-4 and [28]). Moreover, the relative abundance and distribution of these DPR proteins was predictive of toxicity, while the total RNA abundance and RNA foci counts were similar across repeat containing lines, consistent with prior studies [21,28]. While studies in mammalian or human model systems at endogenous expression levels are needed to fully arbitrate the relative roles of RNA and protein in GGGGCC repeat toxicity, our work is consistent with the emerging consensus supporting a significant role of RAN translation in C9ALS/FTD pathogenesis.

In contrast, our finding that the frame in which the C-terminal epitope tag resides was strongly predictive of repeat toxicity is surprising. This finding was quite robust, as we observed this same relationship across multiple different *Drosophila* lines with distinct insertion sites (Supplemental Figure S1) and also when these same constructs were expressed in rodent neurons (Figure 6). The “GFP frame” where the toxicity was greatest would create a fusion with a GP dipeptide repeat protein—the least toxic DPR based on multiple studies using AUG driven expression. Thus, it could be that “out of frame” untagged products are largely driving toxicity while the GFP tag is actually suppressing toxicity derived from certain DPRs. In this context, it is intriguing that the small sequence changes we introduced below the repeat to shift the reading frame resulted in a short disordered 20 amino acid C-terminal tail in the GR reading frame of the repeat in the 1+ (G_4_C_2_)_28_ GFP flies with the most severe phenotypes, with longer C-terminal tails in the GR frame in both the 2+ (G_4_C_2_)_28_ GFP and 0+(G_4_C_2_)_28_GFP flies (Figure 3A, Table 1, and Supplemental Tables S1 and S2). This C-terminal change correlated with both more aggregates of GR DPR in 1+ (G_4_C_2_)_28_ GFP flies and greater nuclear localization of both the GR and other DPRs.

Given that isolated expression of GR dipeptide repeats in the absence of the RNA repeat exhibits marked toxicity in *Drosophila*, rodent and cell based model systems [20,25,53,61,62], changes in its relative solubility, stability or distribution may well influence the toxicity elicited by G_4_C_2_ repeats. In contrast, the relative lack of toxicity associated with the 0+ (G_4_C_2_)_28_ GFP transgene may also be partially explained by a stabilizing effect of a GFP tag on the GA DPR. This GA-GFP fusion product remains largely cytoplasmic and soluble in our assays, which may impede GR mediated toxicity in *Drosophila* [25]. Regardless, our results suggest that the C-terminal region of RAN derived proteins influences their relative toxicity and cellular distribution: a finding with significant implications for design of future model systems at both this and other disease causing repeat expansions.

In rodent neurons, we observed a protective effect when the native C-termini are retained in all three reading frames. Based on studies where we serially enhanced expression of each DPR reading frame by inserting an AUG codon in a strong Kozak context above the GGGGCC repeat element, this protective effect seems to be largely driven by the GR reading frame, as elimination of the C-terminus in GR driven constructs enhanced toxicity while doing so in GA or GP driven constructs had no significant or even a mildly protective effect. Given the propensity for GR DPRs to phase separate[63–65], this native C-terminus may influence the behavior of these peptides when generated in patients. Future studies will be needed to determine if and how such surrounding sequences influence the biochemical behavior of DPRs in general, and GR DPRs in particular.

In summary, our findings demonstrate that the position of the repeat within a transcript as well as its immediate surrounding sequences can have a major impact on its relative toxicity, and on the distribution and abundance of different RAN translated proteins. Future work will be needed to determine the selective effects that these sequence modifications have on the biophysical properties of RAN translated proteins and their exact roles in disease pathogenesis in ALS and FTD. When coupled with recent findings in FXTAS [31] and Spinocerebellar type 36 [66,67], these findings suggest that similar analyses in other repeat disorders are likely warranted.

## List of Abbreviations

ALS: Amyotrophic lateral sclerosis;
FTD: frontotemporal dementia;
RAN translation: repeat-associated non-AUG initiated translation;
DPR: Dipeptide repeats;
UTR: untranslated region;
FXTAS: fragile X-associated tremor/ataxia syndrome;
GA: glycine-alanine;
GP: glycine-proline;
GR: glycine-arginine

## Declarations

### Ethics approval and consent to participate

All mammalian rodent work was approved by the Institutional Animal Care and Use Committee (IACUC) at the University of Michigan.

### Consent for publication

Not applicable.

### Availability of data and material

Not applicable.

### Author Contributions

FH and PKT designed the study. FH, MF, AK, BF, SN, SN, and OA conducted the experiments. FH, AK, and PKT wrote the manuscript. All authors edited the manuscript.

## Acknowledgements

We thank Scott Pletcher, Zhe Han, and Cathy Collins for sharing fly lines. We thank Michael Sutton and the MBNI at Michigan for use of their Confocal Microscope. Reagents generated in the Drosophila Genome research center were supported by NIH Grant 2P40OD010949.

