## Supplemental information for "The carboxyl-termini of RAN translated GGGGCC nucleotide repeat expansions modulate toxicity in models of ALS/FTD"

### Supplemental Table S1: Sequences for $G_4C_2$ Repeat-containing pUAST Vectors

| Name | Sequence | Comments |
|---|---|---|
| GFP-intron | GAATTCGCCACCATGGTGAGCAAGGGCGAGGAGCTGTTCACCGGGGTGGT<br>GCCCATCCTGGTCGAGCTGGACGGCGACGTAAACGGCCACAAGTTCAGCGT<br>GTCCGGCGAGGGCGAGGGCGATGCCACCTACGGCAAGCTGACCCTGAAGT<br>TCATCTGCACCACCGGCAAGCTGCCCGTGCCCTGGCCCACCCTCGTGACCA<br>CCCTGACCTACGGCGTGCAGTGCTTCAGCCGCTACCCCGACCACATGAAGC<br>AGCACGACTTCTTCAAGTCCGCCATGCCCGAAGGCTACGTCCAGGAGCGCA<br>CCATCTTCTTCAAGGACGACGGCAACTACAAGACCCGCGCCGAGGTGAAGT<br>TCGAGGGCGACACCCTGGTGAACCGCATCGAGCTGAAGGgtgagtttccacctagtc<br>caccgtgtgcagcagaaacgagacagagagagagagagagagagagacggggagaaaagtagaagta<br>gtagtagtggtagaaaggaa**gcggccgcctcgag**ggcgcgccactagtgctagcggtacccccttagatctat<br>ctatacccctctctctccctctgtggctctgccccacacggacatggaaatttggaaagacaatcgaccatccatcc<br>gacacacatatctcattcatataccctatatctataacgccaccagGCATCGACTTCAAGGAGGAC<br>GGCAACATCCTGGGGCACAAGCTGGAGTACAACTACAACAGCCACAACGTC<br>TATATCATGGCCGACAAGCAGAAGAACGGCATCAAGGTGAACTTCAAGATCC<br>GCCACAACATCGAGGACGGCAGCGTGCAGCTCGCCGACCACTACCAGCAG<br>AACACCCCCATCGGCGACGGCCCCGTGCTGCTGCCCGACAACCACTACCTG<br>AGCACCCAGTCCGCCCTGAGCAAAGACCCCAACGAGAAGCGCGATCACATG<br>GTCCTGCTGGAGTTCGTGACCGCCGCCGGGATCACTCTCGGCATGGACGA<br>GCTGTACAAGTAATCTAGA | Backbone for all GFP-intronic constructs. Uppercase letters are exons while lowercase are artificial introns. Green highlight indicates interrupted GFP. Red text is the intronic MCS. Restriction sites NotI and XhoI are bolded. |
| NotI-(GGGGCC)$_{3 \text{ or } 21}$-XhoI | **gcggccgc**TACGCATCCCAGTTTGAGACGGGGGCCGGGGCCGGGGCCGGGG<br>CGTGGTCGGGGCGGGCCCGGGGGCGGGCCCGGGGCGGGGCTGCGGTTG<br>CGGTGCCTGCGCCCGCGGCGGCGGAGGCGCAGGCGGTGGCGAGTGGGTG<br>AGTGAGGAGGCGGCATCCTGGCGGGTGGCTGTTTGGGGTTCGGCTGCCGG<br>GAAGAGGCGCGGGTAGAAGCGGGGGCTCTCCTCAGAGCTCGACGCATTTTT<br>ACTTTCCCTCTCATTTCTCTGACCGAAGCTGGGTGTCGGGCTTTCGCCTCTA<br>GCGACTGGTG**ctcgag** | Insert for short $G_4C_2$ repeats. The repeat region is highlighted yellow. The NotI and XhoI sites are red bolded text. |
| NotI-(GGGGCC)$_{28}$-XhoI | **gcggccgc**CCGCAGCCTGTAGCAAGCTCTGGAACTCAGGAGTCGCGCGCTAG<br>GGGCCGGGGCCGGGGCCGGGGCCGGGGCCGGGGCCGGGGCCGGGGCC<br>GGGGCCGGGGCCGGGGCCGGGGCCGGGGCCGGGGCCGGGGCCGGGGC<br>CGGGGCCGGGGCCGGGGCCGGGGCCGGGGCCGGGGCCGGGGCCGGGG<br>CCGGGGCCGGGGCCGGGGCCGGGGCCGGGGCGTGGTCGGGGC*GGGCCC*<br>GGGGGCGGGCCCGGGGCGGGGCTGCGGTTGCGGTGCCTGCGCCCGCGG<br>CGGCGGAGGCGCAGGCGGTGGCGAGTGGGTGAGTGAGGAGGCGGCATCC<br>TGGCGGGTGGCTGTTTGGGGTTCGGCTGCCGGGAAGAGGCGCGGGTAGAA<br>GCGGGGGCTCTCCTCAGAGCTCGACGCATTTTTACTTTCCCTCTCATTTCTC<br>TGACCGAAGCTGGGTGTCGGGCTTTCGCCTCTAGCGACTGGTG**ctcgag** | Insert with 28 $G_4C_2$ repeats is highlighted yellow. The PspOMI site used future self-insertion to make $C_{242}$ and $C_{484}$ is italicized. |
| NotI-(GGGGCC)$_{21}$-PspOMI | **gcggccgc**TACGCATCCCAGTTTGAGACGGGGGCCGGGGCCGGGGCCGGGG<br>CCGGGGCCGGGGCCGGGGCCGGGGCCGGGGCCGGGGCCGGGGCCGGG<br>GCCGGGGCCGGGGCCGGGGCCGGGGCCGGGGCCGGGGCCGGGGCCGG<br>GGCCGGGGCCGGGGCGTGGTC*GGGCCC* | Building block of 21 $G_4C_2$ repeats to generate $C_{49}$, $C_{70}$, $C_{91}$ and $C_{121}$. |

### Supplemental Table S2: Primers used for cloning, In Situ, and qPCR

For C9ORF72 Intronic repeat cloning

| Primer name | Primer sequence |
|---|---|
| NotI-C9 AnchorR | 5'-CCAGCGGCCGCTACGCATCCAGTTTGAGACGGGGGCC-3' |
| XhoI-C9 F | 5'-CCCTCGAGCACCAGTCGCTAGAGGCGAAAGCCCG-3' |
| NotI-C9R | 5'-CCAGCGGCCGCCCGCAGCCTGTAGCAAGC-3' |
| PspOMI-tiling | 5'-AAAGGGCCCGACCACGCCCCGGCC-3' |

For $G_4C_2$ exonic repeat cloning

| Primer name | Primer sequence |
|---|---|
| Adapter1 | 5'-ACCGGTCAGATCTCGAACCGGT-3' |
| Adapter2 | 5'-ACCGGTCAGATCTCGAAACCGGT-3' |

For In situ analysis

| Primer name | Primer sequence |
|---|---|
| Probe | 5'-/5Cy5N/mGmGmCmCmCrCrGrGrCrCrCrCrGr GrCrCrCrCrGrGrCrCrCrCrGmGmCmCmCmC-3' |

For qRT-PCR of GFP in flies

| | |
|---|---|
| GFP unspliced forward | 5'-ACTAGTGCTAGCGGTACCCCTTAGATC-3' |
| GFP spliced forward | 5'-TCTTCTTCAAGGACGACGGCAACTAC-3' |
| GFP reverse (for spliced+unspliced) | 5'-GTACTCCAGCTTGTGCCCCAGGATGT-3' |
| RPL32 forward | 5'-GCCCAGCATACAGGCCCAAG-3' |
| RPL32 reverse | 5'-AAGCGGCGACGCACTCTGTT-3' |

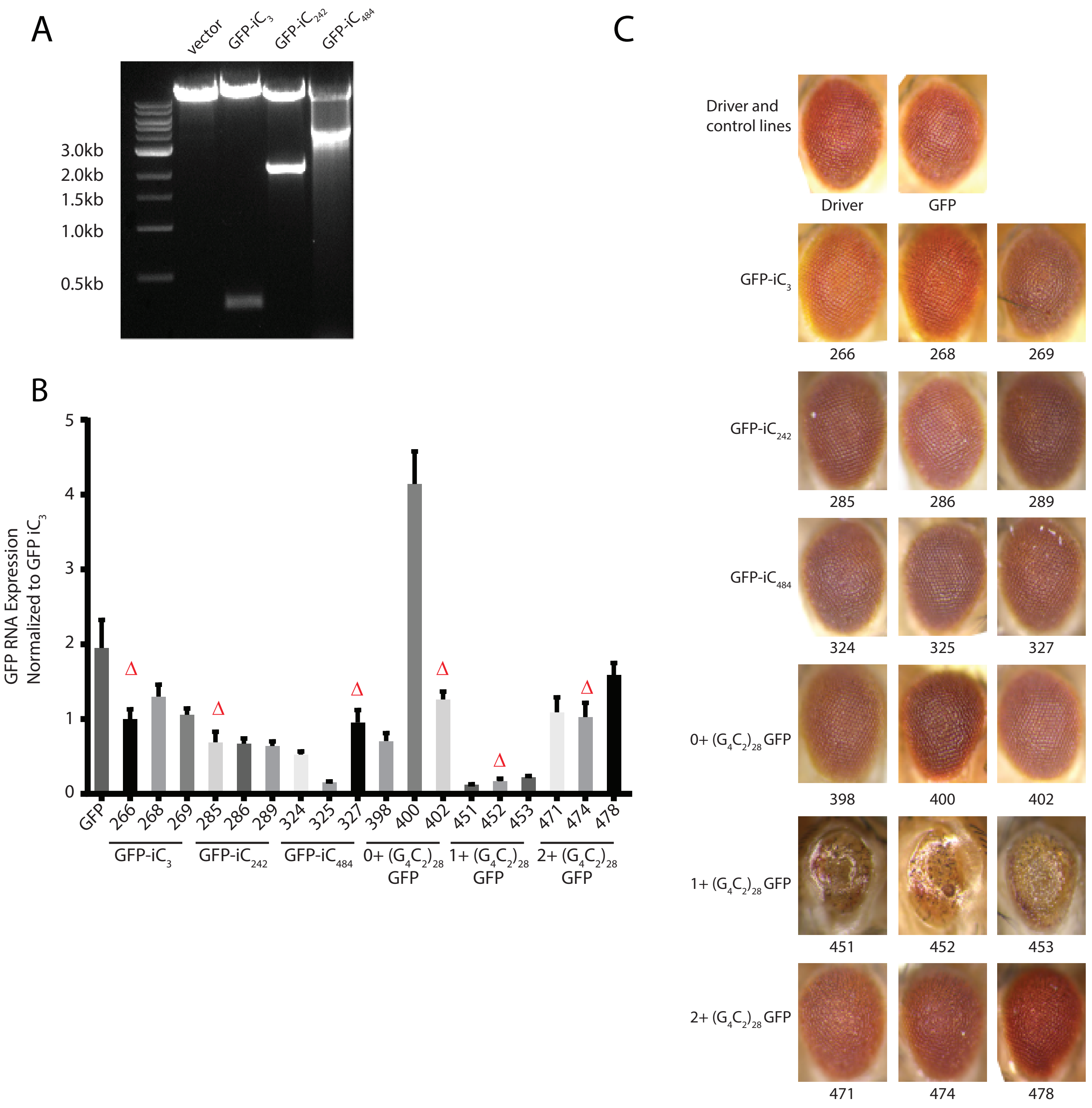
A
vector
GFP-iC3
GFP-iC242
GFP-iC484
3.0kb
2.0kb
1.5kb
1.0kb
0.5kb
B
GFP RNA Expression Normalized to GFP iC3
5
4
3
2
1
0
GFP
266
268
269
285
286
289
324
325
327
398
400
402
451
452
453
471
474
478
GFP-iC3
GFP-iC242
GFP-iC484
0+ (G4C2)28 GFP
1+ (G4C2)28 GFP
2+ (G4C2)28 GFP
C
Driver and control lines
Driver
GFP
GFP-iC3
266
268
269
GFP-iC242
285
286
289
GFP-iC484
324
325
327
0+ (G4C2)28 GFP
398
400
402
1+ (G4C2)28 GFP
451
452
453
2+ (G4C2)28 GFP
471
474
478

**Supplemental Figure S1: Repeat size, RNA expression, and eye phenotypes for intronic and exonic $G_4C_2$ repeat flies.** A) DNA agarose gel validating repeat size of intronic repeat containing vectors. B) GFP mRNA expression from transgenic fly lines. Lines with comparable GFP mRNA levels (Δ) were used for major assays conducted. C) Representative eye images from multiple independent lines for each genotype.

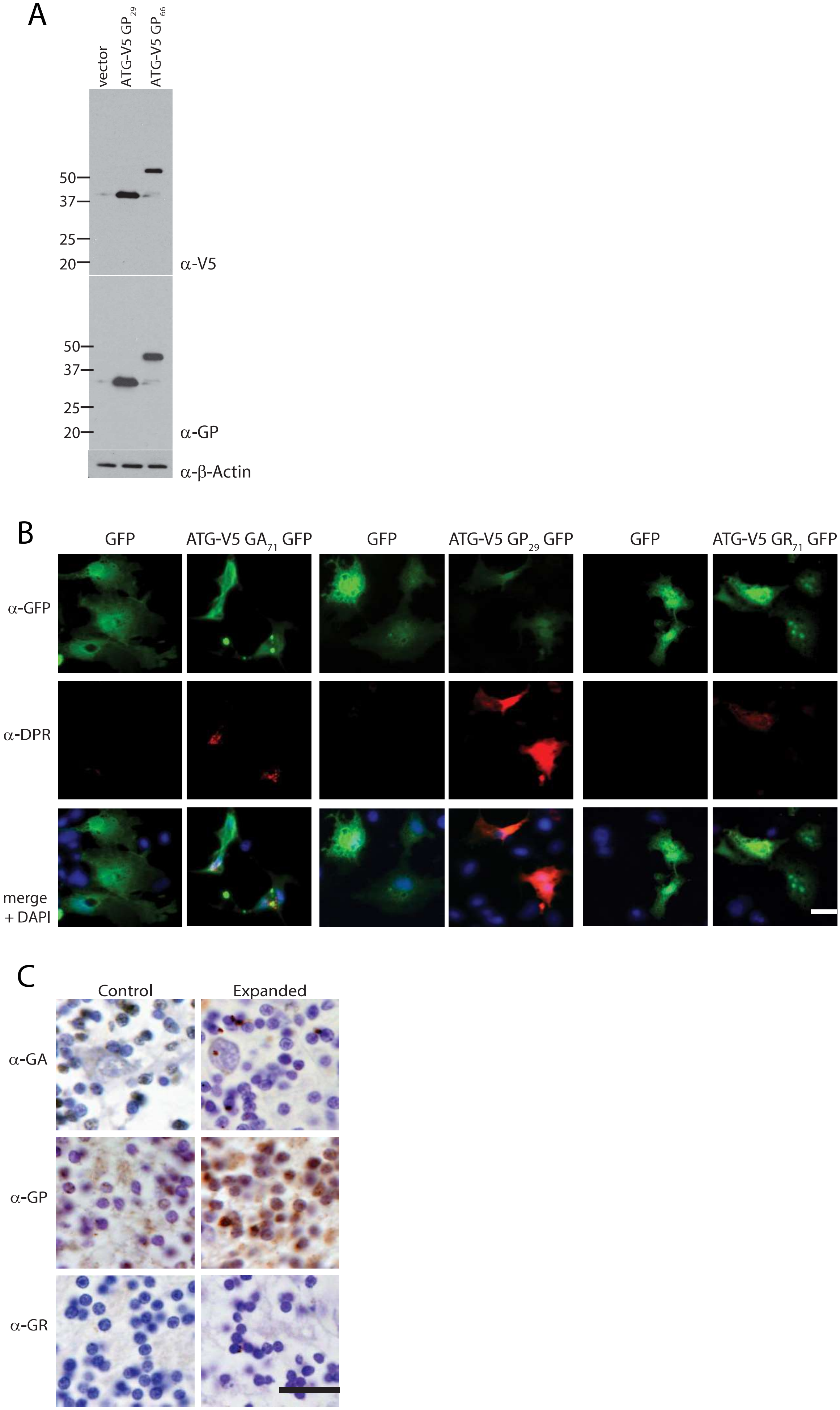
A
vector
ATG-V5 GP29
ATG-V5 GP66
50
37
25
20
α-V5
50
37
25
20
α-GP
α-β-Actin
B
GFP
ATG-V5 GA71 GFP
GFP
ATG-V5 GP29 GFP
GFP
ATG-V5 GR71 GFP
α-GFP
α-DPR
merge + DAPI
C
Control
Expanded
α-GA
α-GP
α-GR

**Supplemental Figure S2: Characterization of dipeptide repeat antibodies.**

Dipeptide repeat (DPR) antibodies specifically recognize $G_4C_2$ containing plasmids by western blot (A) and immunocytochemistry (B). DPR positive aggregates are visible in patient cerebellar tissue (C). Scale bars of B and C = 20μm.
